# What makes our lungs look red?

**DOI:** 10.1101/2021.03.13.435224

**Authors:** Song Huang, Cindia Lopez

**Affiliations:** Epithelix Sàrl, 18 Chemin des Aulx, CH-1228 Plan les Ouates, Geneva, Switzerland

**Keywords:** Stem Cells, Collagen, vWF, Cystic Fibrosis, Pattern Formation, RBC

## Abstract

What makes our lungs look red? This seems to be a naÏve and trivial question. Indeed, all the textbooks tell us that what makes our body red is the presence of blood, or more precisely the red blood cells (RBC). Here we provide some experimental evidence as well arguments to prove that this belief is wrong, or only partially true. In fact, we identified an important population of red cells located outside of the blood vessels as highly compacted clusters, in the connection tissues of the lungs of several species of animals including rat and human being. These red cells possessed each a nucleus, expressing several biomarkers of different cell types such as PF4, vWF, SCF-1R, CD200R, TGF-b, etc… Interestingly, being morphologically heterogeneous, they react collectively to certain stimuli. For example, they aggregated on collagen fibers forming clusters of cells resembling that observed *in vivo*. The red cells may have some features of Hematopoietic Stem Cells, since they were capable of differentiating into other cell types such as alveolar macrophages. In nasal polyps, these cells formed vessel-like structures, confined within a CD31-positive tubes. Upon exposure to toxic chemicals, they formed dense networks, suggesting a possible role in coagulation. Furthermore, the number of these red cells was greatly increased in the lungs of diseased donors, especially in the lungs of CF patients. Instead of being secreted as what happens in normal red cells, vWF proteins were tethered on the cytoplasmic membrane of the red cells isolated from the lungs of CF donors, which may explain at least partially the fibrotic nature of the CF lungs.

Taken together, we conclude that what makes the lungs look red is not the red blood cells, rather a distinct population of red cells, we call them Red Soma Cells (**RSC**). We believe that the discovery and characterization of this important population of cells will have profound theoretic as well as therapeutic implications.

## Introduction

It is generally accepted that what makes our body look red is the presence of Red Blood Cells (RBC), the cellular component of blood. Since RBC contains a large amount of hemoglobin, a red iron-rich protein, therefore it looks red. Due to the presence of a huge number of RBCs, our body also looks red. In this article, we try to prove that this explanation is wrong, or only partially true. To facilitate our demonstration, we want to recall some features of RBCs and the cardiovascular systems [1, 2, 3]:

1. The mammalian red blood cell lacks a nucleus, and it has a biconcave shape.
2. Red blood cells are formed in the red bone marrow from hematopoietic stem cells in a process known as erythropoiesis. Once released into circulation, RBCs are carried by blood flow in the cardiovascular systems to deliver Oxygen and nutrients to all the cells in our body.
3. The cardiovascular systems in the lung has a tree-like hierarchical branching structure. It is not a blind-ending tree, but rather consists of arterial and venous trees that are inter-connected via a capillary bed. A capillary is a small blood vessel from 5 to 10 micrometres (μm) in diameter, and having a wall one endothelial cell thick.
4. The cardiovascular systems are closed, meaning that the blood never leaves the network of blood vessels at normal conditions. If the blood, together with RBCs, leaks from the network of blood vessels, in case of injury or diseases, then the life is in danger.
5. To prevent the blood and RBCs to leave the network of blood vessels upon injury, the activation of coagulation process is needed, initiated by platelet. In other words, RBCs cannot stop moving by itself without the help of platelets.

In this article, we will show that in the lungs there exists an important population of cells, despite of their red color and RBC-like morphology, it is a distinct cell type. They are located outside of the blood vessels, more precisely in the connective tissues of the lungs of several species of animals, such as Rat and Human. These red cells possess each a nucleus, expressing numerous biomarkers of hematopoietic cells, such as PF4, vWF, SCF-1R, CD200R, TGF-β, etc… Interestingly, being morphologically heterogeneous, they react collectively to certain stimuli. For example, they attach and aggregate on collagen fibers, showing a collective behavior. In nasal polyps, these cells formed vessel-like structures, confined within a CD31-positive tubes. Upon exposure to toxic chemicals, they form dense networks, suggesting a possible role in coagulation. Furthermore, the number of these red cells was greatly increased in the lungs of diseased donors, especially in the lungs of Cystic Fibrosis (CF) patients. Instead of being secreted as what happens in normal red cells, vWF proteins are tethered on the cytoplasmic membrane of the red cells isolated from the lungs of CF donors, which may explain at least partially the fibrotic nature of the CF lungs.

Taken together, we conclude that what makes the lungs look red is not the red blood cells, rather a distinct population of red cells, we call them Red Soma Cells (**RSC**). We believe that the discovery of this important population of cells will have profound theoretic and therapeutic implications.

## Materials and Methods

### 1. Chemicals

37% Forlmaldehyde, Poly (4-styrenesulfonic acid), Glycerol, Methanol, Trypsin were purchased from Sigma-Aldrich. Blue Ink, from Schrebwaren GmbH. Type I Collagen, Collagenase and Elastase, from Worthington, USA. Pronase, from Roche diagnostics GmbH. All the culture media, such as RPMI culture medium, DMEM culture medium, were purchased from Life Science (Gibco).

### 2. Lung biopsies, isolation and culture of RSC cells

a. The lungs or lung biopsies were obtained from patients having lung transplantation or polypectomy. All experimental procedures were explained in full, and all subjects provided informed consent. The study was conducted according to the declaration of Helsinki on biomedical research (Hong Kong amendment, 1989), and received approval from local ethics commission.
b. Isolation and purification of RSC cells A large number of RSC cells exist in the lungs, especially in the lung of donors with inflammatory diseases. Therefore, RSC cells could be obtained easily from minced lung parenchyma. If needed, RSC cells can be further purified by Ficoll gradient centrifugation to get rid of contaminating cells such as alveolar macrophages, by following the same protocol for isolating the Peripheral Blood Monocyte Cells (PBMC). Macrophages are in suspension and RSC cells at the bottom of tube as cell pellet. RSC cells are cultured in RPMI supplemented with 10% fetal bovine serum and antibiotics (RPMI^++^).
c. Enzymatic digestion of the lung parenchyma The lung tissue was dissected into small pieces about 5 mm^3^. Then, the minced tissues (about 10 g) were placed into 50 mL conical tubes, filled washing buffer (Dulbecco’s Phosphate Buffered Saline (DPBS) containing 100U/mL each of penicillin and streptomycin). Tubes were inverted to wash the tissue and the supernatant discarded, repeating for three times. To remove as much as possible the liquid, the minced and washed tissues were filtered through a layer of sterile gauze. The tissues on the Gauze were distributed to 15 ml of Falcon tube, 1.5 g of tissues per tube. Then, the tissues were digested with 5 ml of enzymes. All the enzymes were prepared in PBS without Ca^++^ and Mg^++^. The concentration of the enzyme: 0.01% (w/v) elastase, 0.5% (w/v) collagenase, 0.25% (w/v) trypsin, 1.5 mg/ml Pronase. The digestion was carried out in a Water bath at 37°C, for 2 hours. After enzymatic digestion, 8 ml of the PBS were added into each tube, then filtered again through a sterile Gauze. If needed, pass the filtrates through a 10 µm cell strainer. Collect the cells by centrifugation.

### 3. Ink staining of the rat lungs

The rat lungs were dissected out together with the heart, then about 3 ml of a blue ink was injected into the right ventricle of the rat heart. As expected, the ink entered into the lungs and stained the blood vessels. After ink injection, the whole rat lungs were fixed in 4% Formaldehyde for 24 hour at 4°C. The lung samples were observed after being cleared by increasing concentrations of Gylcerol diluted in 0.5% KOH solution (1/4, 1/2, 3/4, and pure Glycerol). Or the lungs were cut into slices of 100 µm of thickness with a McILWAIN Tissue chopper (made by The Mickle Laboratory Engineering CO, LTD, commercialized by World Precision Instruments, Hertfordshire, UK).

### 4. Immuno-cytochemistry

Paraffin sections of the lung tissues were performed according to the standard procedures. The lung tissues were fixed in 4 % Formaldehyde. RSC cells were fixed first with a solution of 4% Paraformaldehyde and 1% Glutaraldehyde in 0.1M Phosphate buffer at pH 7.3, at room temperature for 1 hour; then fixed again with cold Methanol for 30 min. Methanol will make the cells to stick to the bottom of the plates, facilitating the following steps of Immuno-cytochemistry. The primary antibodies used were anti-vWF (1:50, Dako, GA527); anti-TGF-β (1:25, Thermofisher Scientific, MA5-23795); anti-CD31 (1:50, Dako, GA610); anti-CSF-1R (1:25, Invitrogen, PA5-14569); anti-PF4 (1:50, Sigma, SAB4301756); anti-CD200R (1:25, Merck, HPA046404). Four secondary antibodies were purchased Molecular Probes, Thermofisher Scientific: the goat anti-mouse IgG Alexa fluor 488, the goat anti-mouse IgG Alexa fluor 568, the goat anti-rabbit Alexa fluor 488 and the goat anti-rabbit Alexa Fluor 568. The antibodies are incubated for one hour at room temperature or overnight at 4°C. After washing, the secondary antibody was applied on samples. The fluorescent signal of each protein was visualized and digitalized using a Leica DMIRE2 Fluorescence Microscope, and ImageProplus software.

### Results

#### 1. Presence of red colored cells in the lung biopsies

While working with lung biopsies, be it nasal polyps, bronchi, or parenchyma, we noticed that along with the biopsies came a large number of small, red cells that we initially considered as red blood cells. However, upon closer examination, these red cells didn’t appear to be confined within the blood vessels, instead, they seemed to be trapped in the connective tissues (Figure 1 a, b). Under the contrast microscope, the red cells were indeed intermingled with collagen fibers (Figure 1 c, d). Consistent with these observations, these red cells could be isolated more efficiently from the same amount of minced human lung parenchyma with collagenase than with other enzymes, such as elastase, pronase or trypsin (Figure e, f). Intriguingly, among the RBC-like cells, some red cells had an appearance of monocytes, or neutrophils (Figure 1 g, h). Therefore, it was possible that these red cells represent a distinct cell population, instead of being red blood cells escaped from the blood vessels, clotted on lung biopsies.

**Figure 1:**
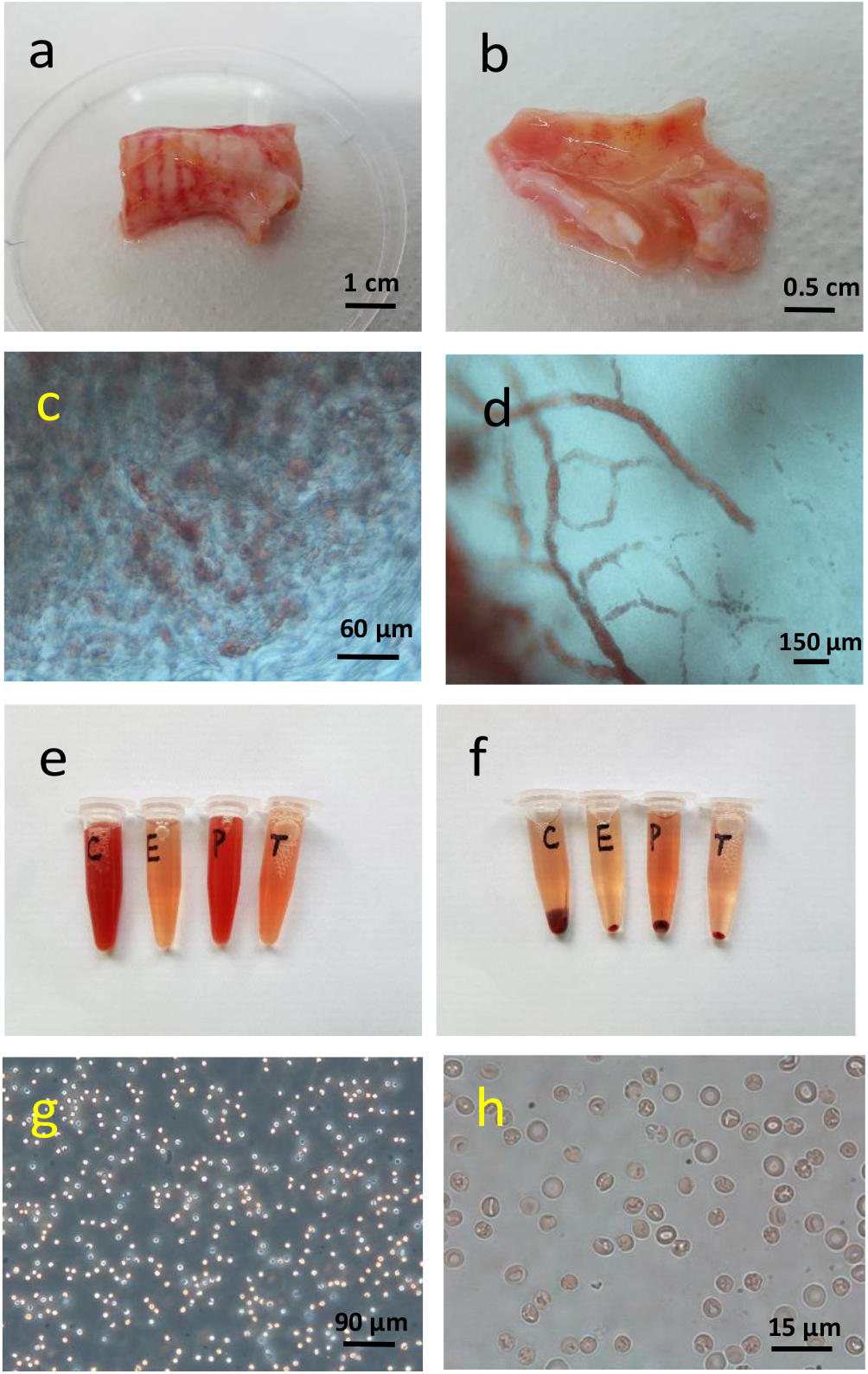
The red cells were present in lung biopsies. (a) and (b), segments of human bronchus, showing the stripes of red cells on both sides of the bronchial tubes. (c), a microscopic image of a piece of connective tissues surrounding the bronchial tube, showing the red cells intermingled with the collagen fibers. (d), the red cells wrapped in collagen fibers taken from human nasal polyps. (e) and (f), the red cells isolated from 1.5 g of lung parenchyma, after 2 hours of enzymatic digestions (C, collagenase; E, Elastase; P, Pronase; T, Trypsin); (e) before centrifugation; (f), after centrifugation. (g) and (h), Phase contrast microscope images showing the morphologies of the red cells.

#### 2. These red cells are located outside of the blood vessels in the rat and human lungs

Since the biopsies have been surgically removed from the donor’s body, it was possible that the biopsies were “contaminated” by blood and red blood cells during the surgical operations. In order to avoid such scenario, we performed an experiment with intact organs devoid of any mechanical injury of the lungs (no incision nor cut). The rat lungs were dissected out together with the heart, then a blue ink was injected into the right ventricle of the rat heart. As expected, the ink entered into the lungs and stained the blood vessels (Figure 2 a). The color of the control rat lungs, without ink staining, appeared more uniform (Figure 2b). Under the contrast phase microscope, the arterioles (Figure 2c) and capillary bed (Figure 2d) were both stained blue. What is evident was the presence of the red colored cells outside of the blue vessels. Since the injected ink was contained inside of the blood vessels, this means that the blood vessels were intact and sealed, thus the red blood cells had no chance to get out of the circulation, and to “contaminate” the surrounding lung tissues. In other words, the red cells in Figure 2 c and d could not be RBCs escaped from the blood vessels, rather they may be true resident pulmonary cells.

**Figure 2:**
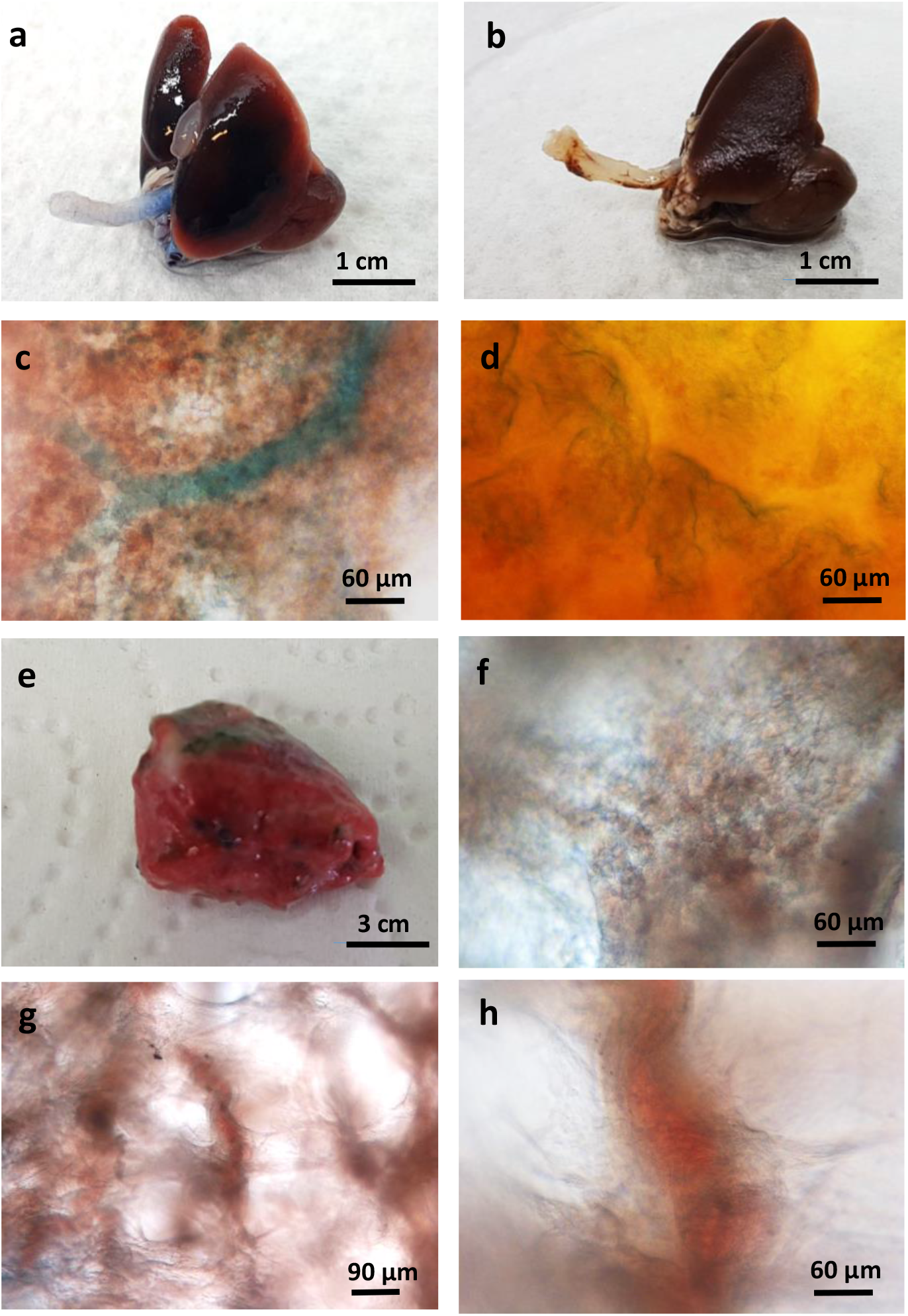
The red cells were located outside of the blood vessels. (a), the rat lungs were stained with a blue ink by injection of the ink into the right ventricle of the heart. (b), the control rat lungs without ink staining. (c, d), the slice of the ink stained lungs observed under the contrast microscope after tissue-clearing with Glycerol. (e), a piece of human lung parenchyma from a diseased donor, showing the dark red color. (f), a slice of the human lungs observed under the contrast microscope after tissue-clearing with Glycerol, showing the red colored cells on the alveolar wall. The capillaries appeared blueish on the picture. (g) and (h), Phase contrast microscope images showing the presence of highly compact clusters of red cells in the septa of the alveoli.

In fact, hundreds of similar experiments are performed almost daily in numerous hospitals around the world! The human lungs that we received from our collection centers were donated initially for transplantation, but due to diverse reasons, some lungs were not suitable for this purpose, thus they were redirected for research. In order to preserve the quality of the organs, according to a standard procedure [4], the lungs were flushed with a chilled acellular preservation solution through the pulmonary artery. In other words, the red blood cells along with the blood have been chased out of the lungs before being surgically removed from the donor’s body. Despite of lacking blood and RBCs, the lungs that we received remained red (Figure 2 e). Under the microscope, the red cells showed a similar structure as that found in the rat lungs: they formed an integral part of the alveolar walls (Figure 2 f), or densely packed in alveolar septa (Figure 2 g, h). Even in absence of ink staining, the capillary bed looked blueish and could be easily recognized (Figure 2 f). In the majority of the cases, the lungs that we requested and received came from healthy donors, at least no documented lung diseases. Therefore, it is unlikely that these donors had massive internal hemorrhage in the lungs before being prepared for transplantation.

Taken together, we conclude that there exists an important population of red cells in the lung, distinct from the red blood cells, which are located outside of the blood vessels, most likely within the connective tissues rich in collagen fibers. We name this population of resident red pulmonary cells, **Red Soma Cells**.

#### 3. The Red Soma Cells could attach, aggregate and form patterns on collagen fibers *in vitro*

Consistent with our observation *in situ*, the Red Soma Cells were capable to attach and to aggregate on collagen fibers *in vitro*. By drying a solution of 100 µg/ml collagen dissolved in PBS, we were able to create some random patterns formed by salt crystals at the bottom of a 24 well plate (Figure 3a). Interestingly, when seeded onto the patterned wells, the Red Soma Cells replicated the patterns formed by salt crystals and collagen fibers (Figure 3b, c). Therefore, this was a collagen fiber-induced collective and self-organizing behavior. What was surprising, despite of the obvious morphological difference among these red cells, almost all the cells in the picture were capable of attaching to the collagen fibers and forming aggregates (Figure 3c). In absence of collagen and collagen fibers, the Red Soma Cells were homogeneously distributed at the bottom of the wells, despite of a high cell density (Figure 3d). Based on this criteria, these cells could be considered as a single cell type.

**Figure 3:**
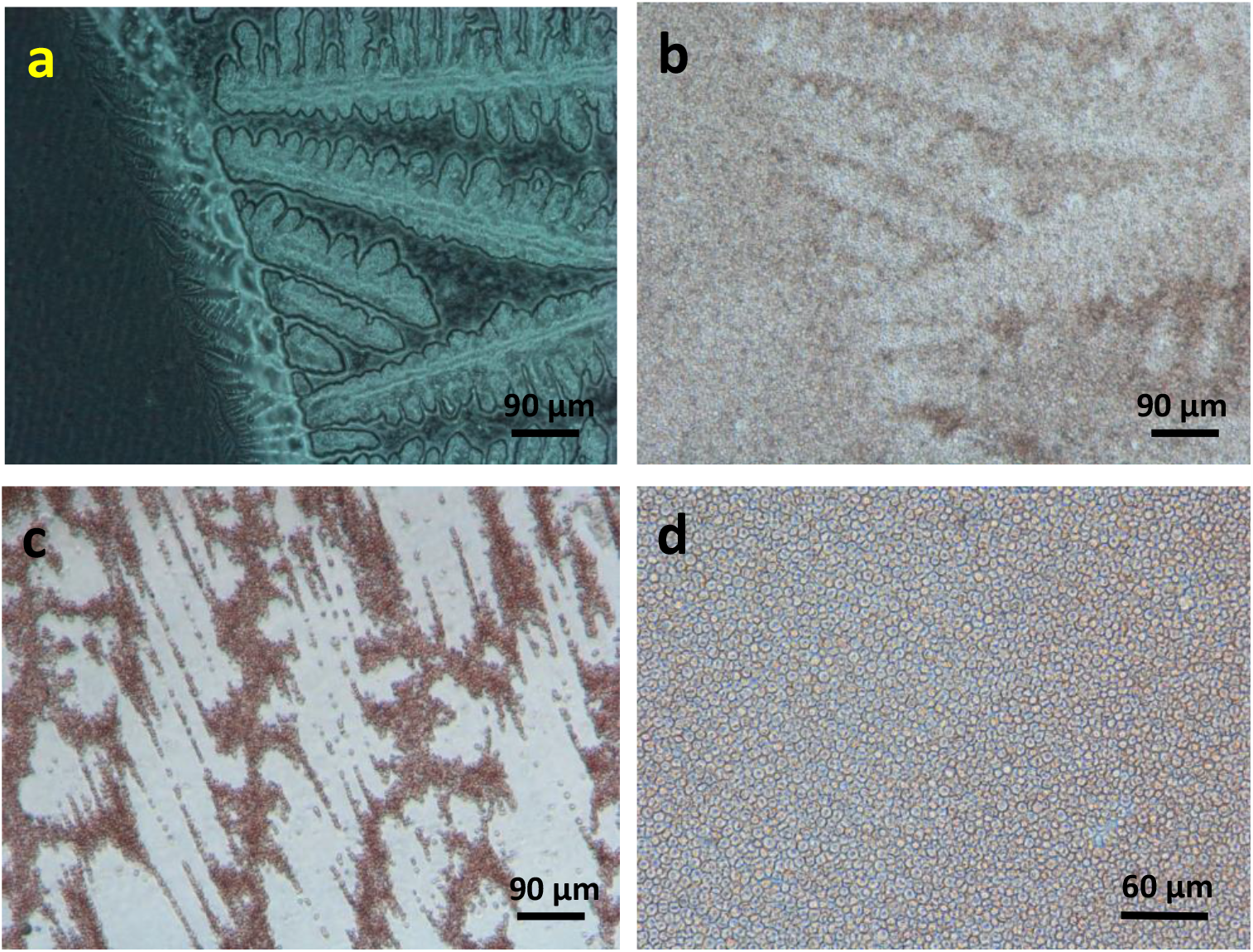
The red cells showed affinity to collagen and formed patterns on collagen fibers. (a) By drying a solution of 100 µg/ml collagen dissolved in PBS, some random patterns **w**ere formed by salt crystals at the bottom of a 24 well plate, especially at the periphery of the wells. (b). the Red Soma Cells replicated the patterns formed by salt crystals and collagen fibers. (c). the Red Soma Cells attached to collagen fibers, forming clusters of cells. (d), in absence of collagen fibers, the cells were evenly distributed at the bottom of the well, despite of a high cell density.

#### 4. The Red Soma Cells expressed numerous biomarkers of hematopoietic cells

Due to their resemblance to RBCs, these Red Soma Cells may be some kind of hematopoietic cells. First of all, the Red Soma Cells contained a nucleus since they were stained positive with a DNA dye, Dapi (Figure 4 a, b). This result alone would distinguish these cells, without any ambiguity, from the Red Blood Cells that lack a nucleus. Using specific antibodies, we showed that these Red Soma Cells expressed a variety of biomarkers, such as vWF, PF4, CSF-1R, CD200R, etc… The positive signal of vWF staining appeared as dots, presumably a structure similar to Weibel-Palade bodies usually found in endothelial cells ((Figure 5 c). Interestingly, not all the cells were stained positive for vWF, presumably the vWF was already secreted. It has been known that an induced secretion of VWF is associated with the loss of Weibel-Palade bodies [5]. PF4 was also detected in RSC cells (Figure 5 d). As expected, CSF-1 receptor was located on the cytoplasmic membrane (Figure 5 e). Interestingly, Red Soma Cells also expressed biomarkers of alveolar macrophages, such as CD200R (Figure 5 f), among others. We have looked only a limited number of biomarkers, but it was already quite impressive, considering the small size of these red cells. Moreover, the nucleus was almost as big as the cell itself.

**Figure 4:**
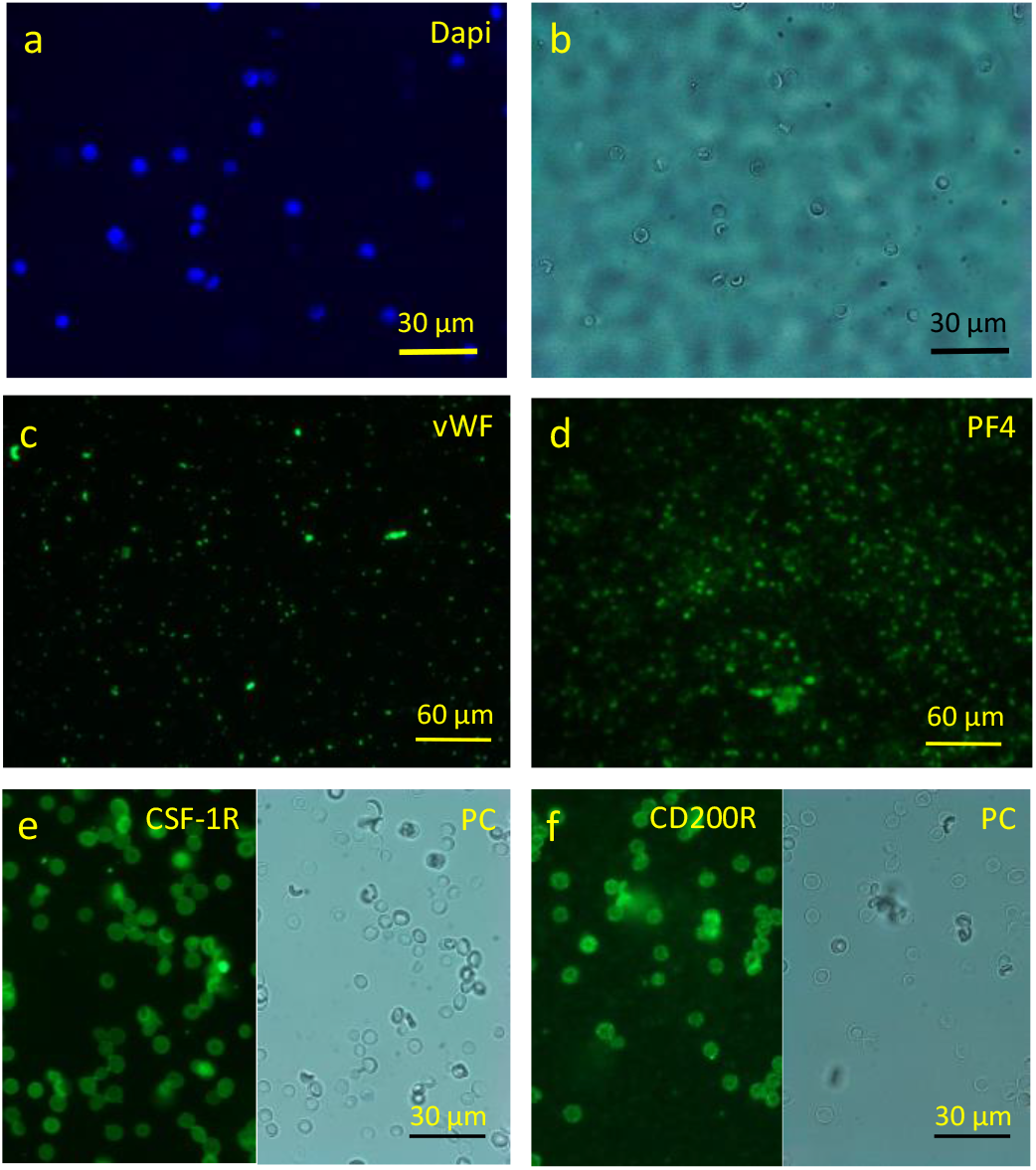
The Red Soma cells expressed a variety of biomarkers. RSCs stained with a DNA dye, Dapi. (b). Phase contrast (PC) image of the same cells in (a). (c). Immuno-staining of the Red Soma Cells with anti-vWF antibody. (d). Immuno-staining of the Red Soma Cells with anti-PF4 antibody. (e). Immuno-staining of the Red Soma Cells with anti-CSR-1R antibody. (f). Immuno-staining of the Red Soma Cells with anti-CD200R antibody.

**Figure 5:**
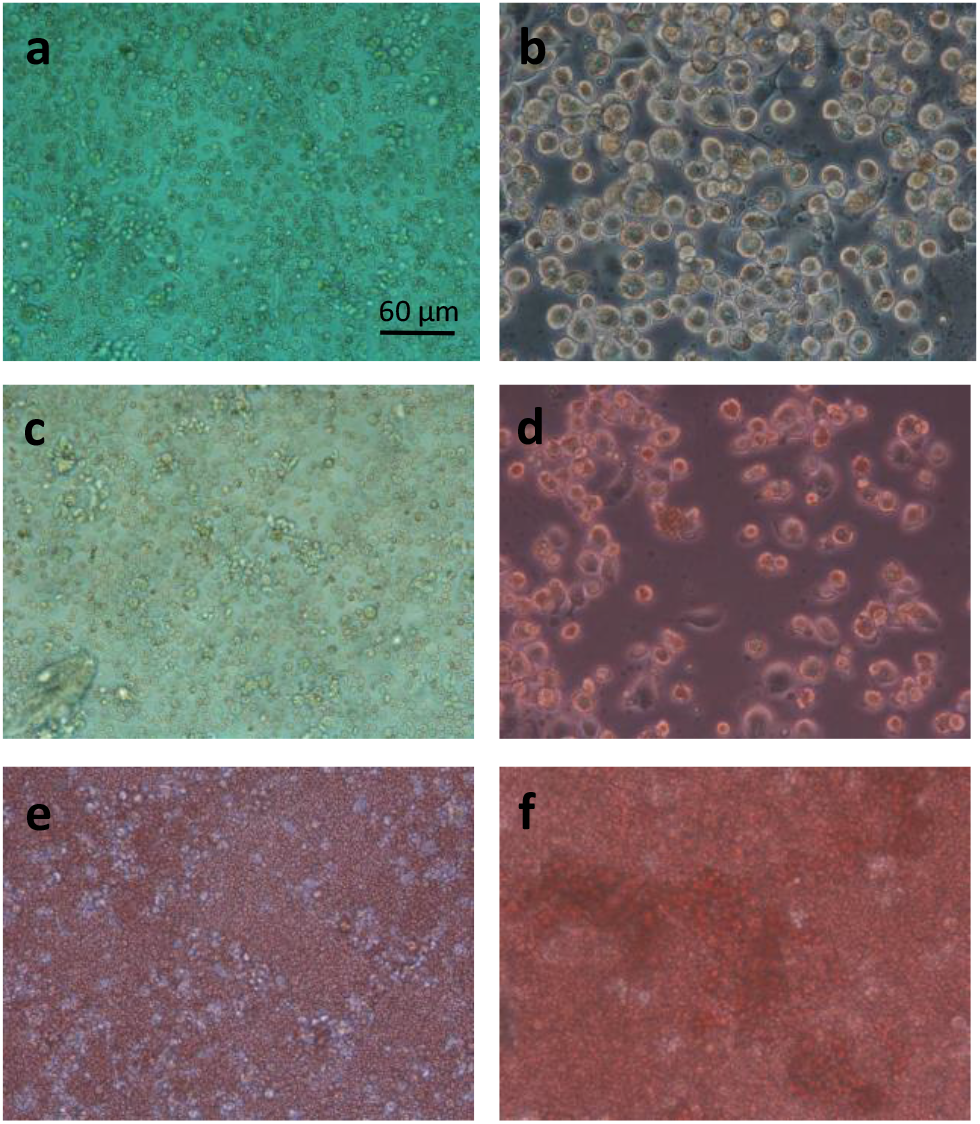
Generation of macrophages from the rat Red Soma cells in culture. Rat RSC cells were cultured in media complemented with 10% Fetal Bovine Serum (FBS): (a, b), RPMI culture medium; (c. d) DMEM; or (e, f), in a serum-free culture medium (MucilAir culture medium, Epithelix). (a, c, e), pictured taken at D1. (b, d, f), pictured taken at D7. All the pictures were taken under the same setting of imaging system (bar = 60 µm).

#### 5. Possible functions of RSC cells

Judging by their large number and the expression of numerous biomarkers, as well as their collective behavior on collagen fibers, we speculated that RSC cells must have some important functions in the lungs. We would like to show below some possible functions of RSC cells in the lungs, such as generation of alveolar macrophages, angiogenesis, coagulation, and CF pathogenesis. Due the limitation of the space, it was not possible to explore all these aspects in depth in one article. Indeed, each phenomenon described below would constitute a research subject on its own. However, the evidence presented here would be sufficient to demonstrate the versatility, the plasticity, the flexibility as well as the unity of RSC cells.

#### 5.1. Generation of Macrophages

Since RSC cells expressed several biomarkers of alveolar macrophages such as CD200R, TGF-β, RSC cells may have the potential to differentiate into macrophages. In fact, a large number of RSC cells could be co-purified with the alveolar macrophages. In lung sections or slices, RSC cells were often found in the vicinity of alveolar macrophages. Nevertheless, we had difficulty to obtain sufficient number of mature macrophages from the human RSC cells at our current culture conditions. Proportionally, the number of macrophage-like cells was relatively low. Moreover, the macrophages entered into apoptosis after several days of culture. However, this was not the case for RSC cells from the rat lungs. Morphologically mature macrophages could easily be observed after one week of culture *in vitro:* RSC cells at Day 1(Figure 5 a, c, e) cultured in different media; RSC cells at Day 7 (Figure 5 b, d, and f). This differentiation process seemed to be serum-dependent, because a large number of mature macrophages were present in the culture media supplemented with 10% fetal bovine serum (Figure 5 b, d), few in a synthetic, serum-free culture medium (figure 5 f).

#### 5.2. Angiogenesis

Nasal polyps are noncancerous growths within the nose or sinuses [6]. They are commonly associated with conditions that cause long term inflammation of the sinuses. This includes chronic rhinosinusitis, asthma, aspirin sensitivity, and cystic fibrosis [7]. Interestingly, vessel-like structures could be frequently observed in nasal polyps (Figure 6a), suggesting that there may have de nouveau formation of blood vessels, namely angiogenesis. Using a specific biomarker of the endothelial cells, CD31, we showed that these vessel-like structures were indeed stained positive with anti-CD31 antibody (Figure 6b). The cells inside the vessels were labeled by anti-vWF antibody (Figure 6, c and d). Double IF staining showed that these two biomarkers were closely related, vWF appeared as small dots, and CD31 as fibers (Figure 6, e and f). These results suggested a potential role of RSC cells in angiogenesis. Like a root bud of a plant [8], RSC cells may “drill through” the collagen fibers to form vessel-like structure. Thus, even though most of RSC cells resided outside of the cardiovascular system in the adult, they may also be inside of the blood vessels during certain stages of the embryonic development of the lung or at certain disease conditions.

**Figure 6:**
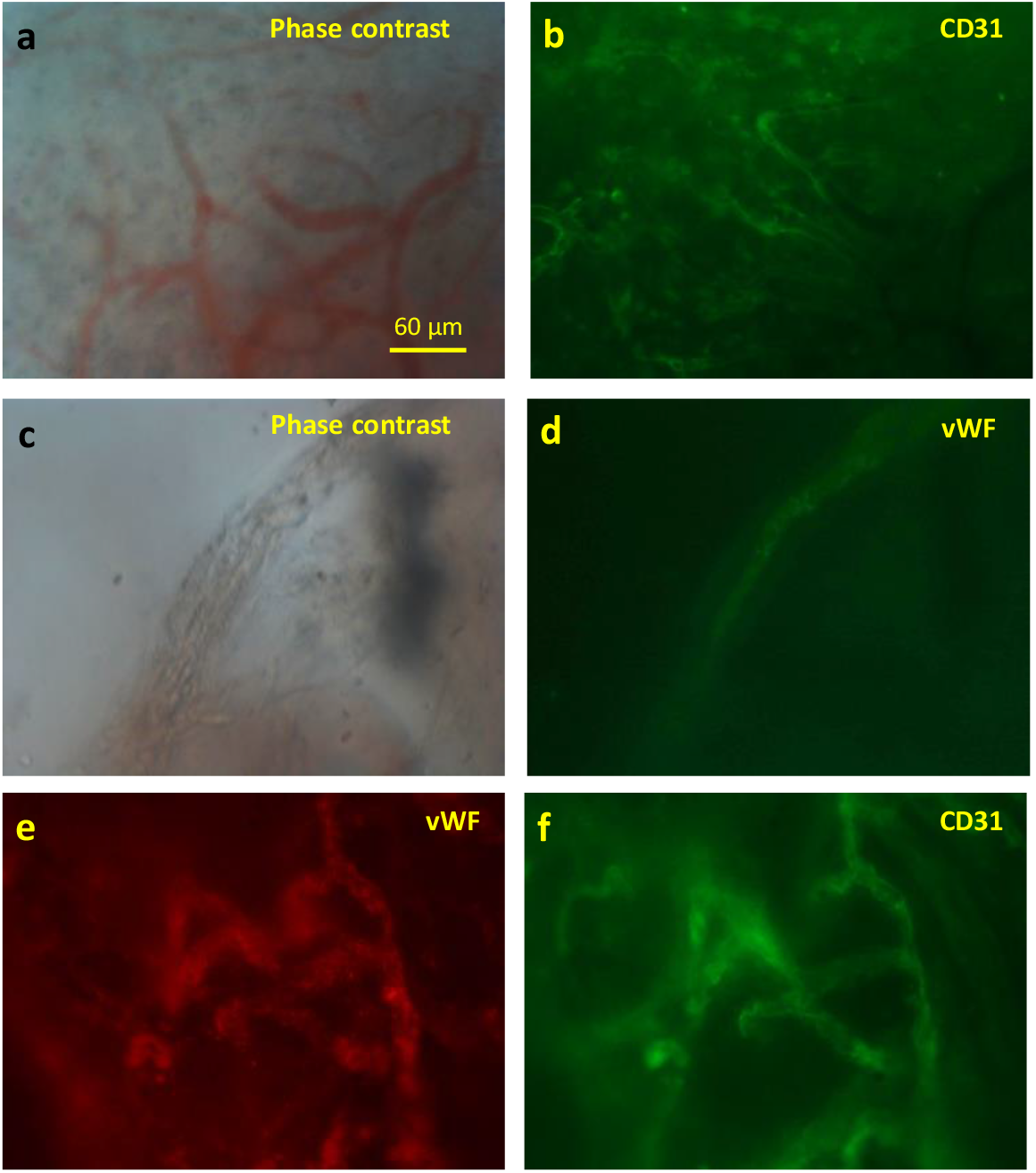
Possible involvement of RSC cells in angiogenesis. (a), Vessel-like structures in biopsies of nasal polyps. (b, f), In situ immuno-staining with anti-CD31 antibody. (c), Phase contrast image of (d). (d, e), *In situ* immuno-staining with anti-vWF antibody. All the pictures were taken under the same setting (bar = 60 µm).

#### 5.3. Coagulation

Since RSC cells expressed two markers of megakaryocytes and platelets, PF4 and vWF, we wondered if RSC cells could have a role in coagulation. To this end, we treated these cells with some toxic chemicals. One of them, Poly (4-styrenesulfonic acid) (PSS), was able to activate these cells in a dose-dependent manner. At low concentrations (≤ 0.5%) of PSS, most of RSC cells kept their morphology, but some closely related cells formed cell-stacks (Figure 7a). However, at higher concentrations (> 1%), the RSC cells rapidly morphed into filaments, forming interconnected networks (Figure 7b, c, d, e). Since, the cell density was a critical factor for network formation, we placed a dense monolayer of RSC cells on coverslip in 400 µl of culture medium, and then we added the same volume of PSS at a given concentration. The fact to mix two solution generated a PSS gradient and a dynamic flow of the liquid. This process may have caused the formation of different patterns and networks depicted in Figure 7. Like the pattern formation of RSC cells on collagen fibers, the network formation could also be an active and dynamic process. It was clearly a collective and collaborative behavior. It seemed that RSC cells could sense the danger signal in the environment and respond accordingly. Indeed, if we looked carefully the structure and composition of the network, we could distinguish at least two modes of actions: colloidal mode and polymer mode. In Figure 7b, the cells remained globular but anisotropic. They formed an interconnected network by joining the adjacent RSC cells. However, at certain conditions, the whole cell was transformed into a long filament (Figure 7f), then the pattern formation was based on percolation of the polymers (Figure 7 c, d). With the help of the dynamic force of liquid flow, the long fibers could form at long distance (Figure 7e). Thus, this experiment demonstrated the intrinsic ability of RSC cells to morph into fibers and to form networks. This could be an important step in coagulation of blood.

**Figure 7:**
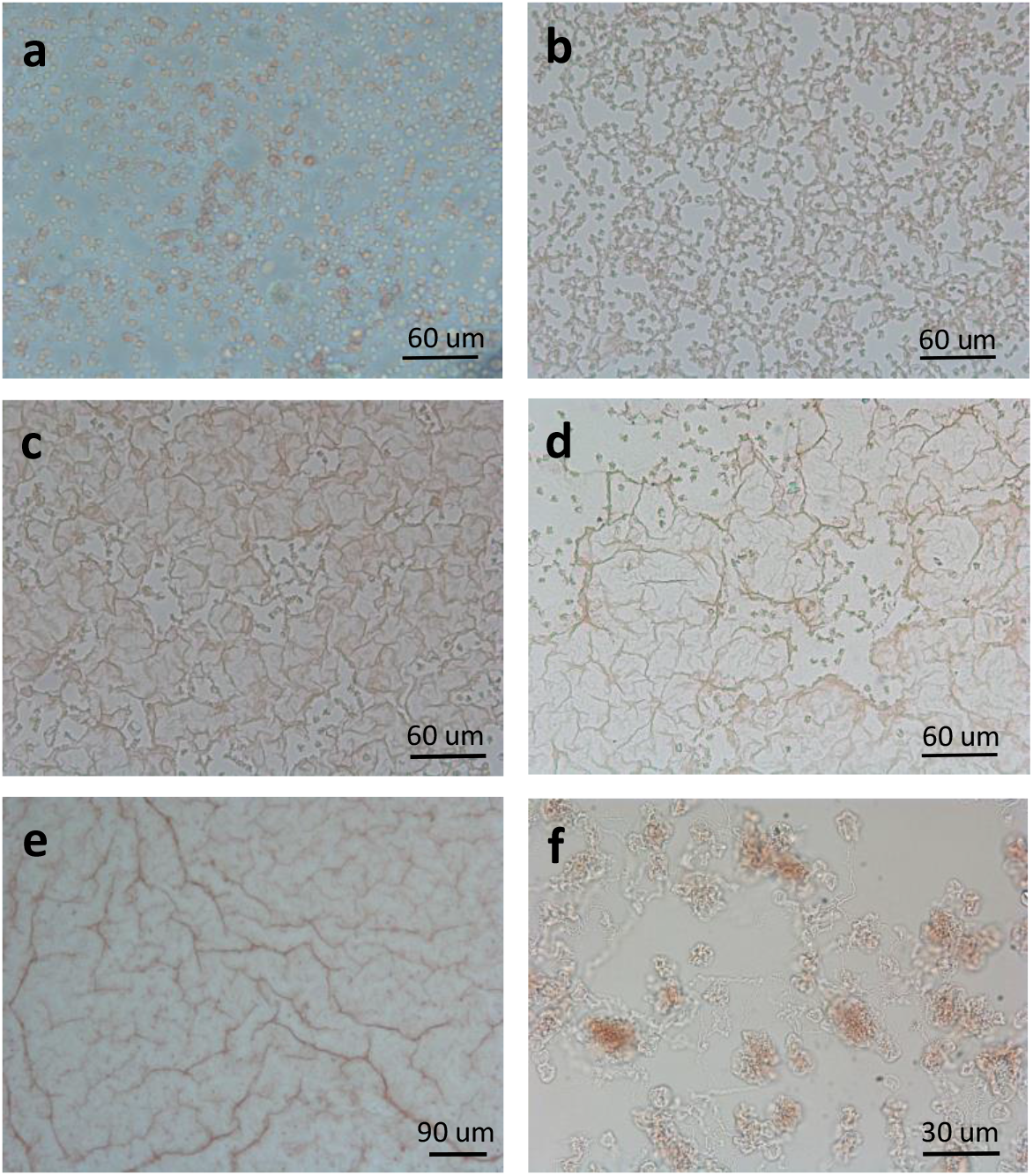
Possible function of RSC cells in coagulation. (a), RSC cells treated with 0.5% PSS solution; (b, c, d, e, f), with 2 % PSS solution.

#### 6. Possible involvement of RSC cells in the pathogenesis of Cystic Fibrosis

##### 6.1. Abnormal appearance and structure of Cystic fibrosis (CF) lungs

CF is the most frequent genetic diseases with an autosomal recessive feature. It is caused by mutations in the CFTR gene [9]. CFTR dysfunction affects many organs with complications including sinusitis, diabetes mellitus, bowel obstruction, hepatobiliary disease, hyponatremic dehydration, and infertility [10]. However, the major cause of morbidity and mortality is the progressive bronchiectasis and ultimately respiratory failure, due to chronic infection and inflammation of the airways in patients with CF [11, 12]. The lung biopsies from the CF donors did show obvious abnormalities: often the bronchial tubes had “bloody” appearance, and surrounded by lumps of red cells (Figure 8a, CF). On the histological sections, certain regions of lung parenchyma from CF donors were filled with cells, and the alveolar structures were destroyed (Figure 8b, c). Clearly, the lungs of CF donors were fibrotic in nature. This is unlikely an experimental artefact because, processed the same way, the alveolar structures of the non CF lungs remained intact and looked normal (Figure 8b, d).

**Figure 8.**
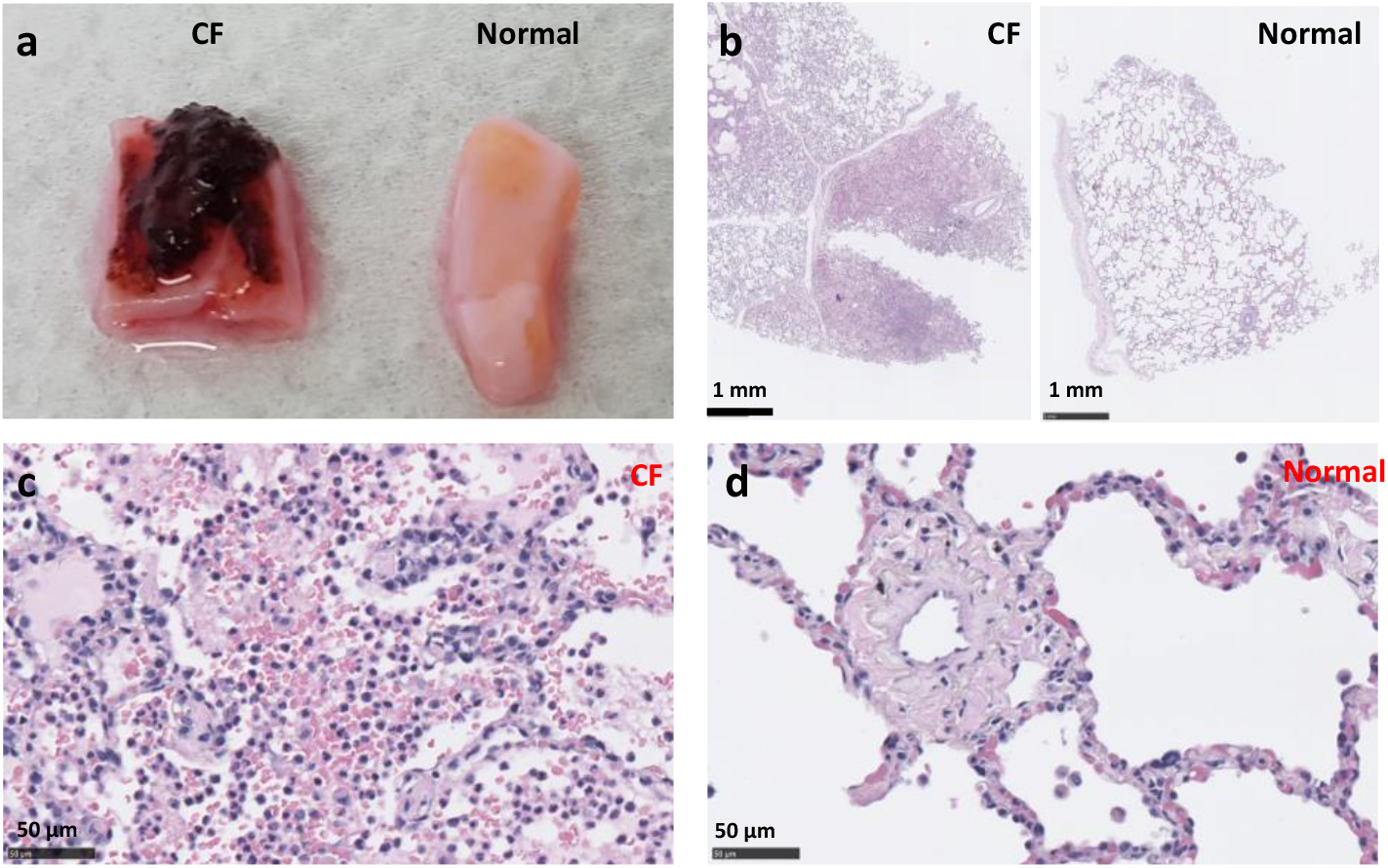
Abnormal structures of the CF lungs. (a), the bronchial tube from CF donors were covered with lumps of red cells. (b, c), On sections of CF lung parenchyma, fibrotic tissues could be seen. (d), Picture depicts the structure of alveoli in non-CF lung.

##### 6.2. Mislocalization of vWF proteins in CF RSC cells

TGF-β is a key growth factor in promoting lung fibrosis [13, 14]. Thus we stained the lungs sections, from CF and non-CF donors, with anti-TGF-β and vWF antibodies. As expected, TGF-β was expressed in both CF and non-CF lungs, obviously in the connective tissues (Figure 9 b, e).

**Figure 9.**
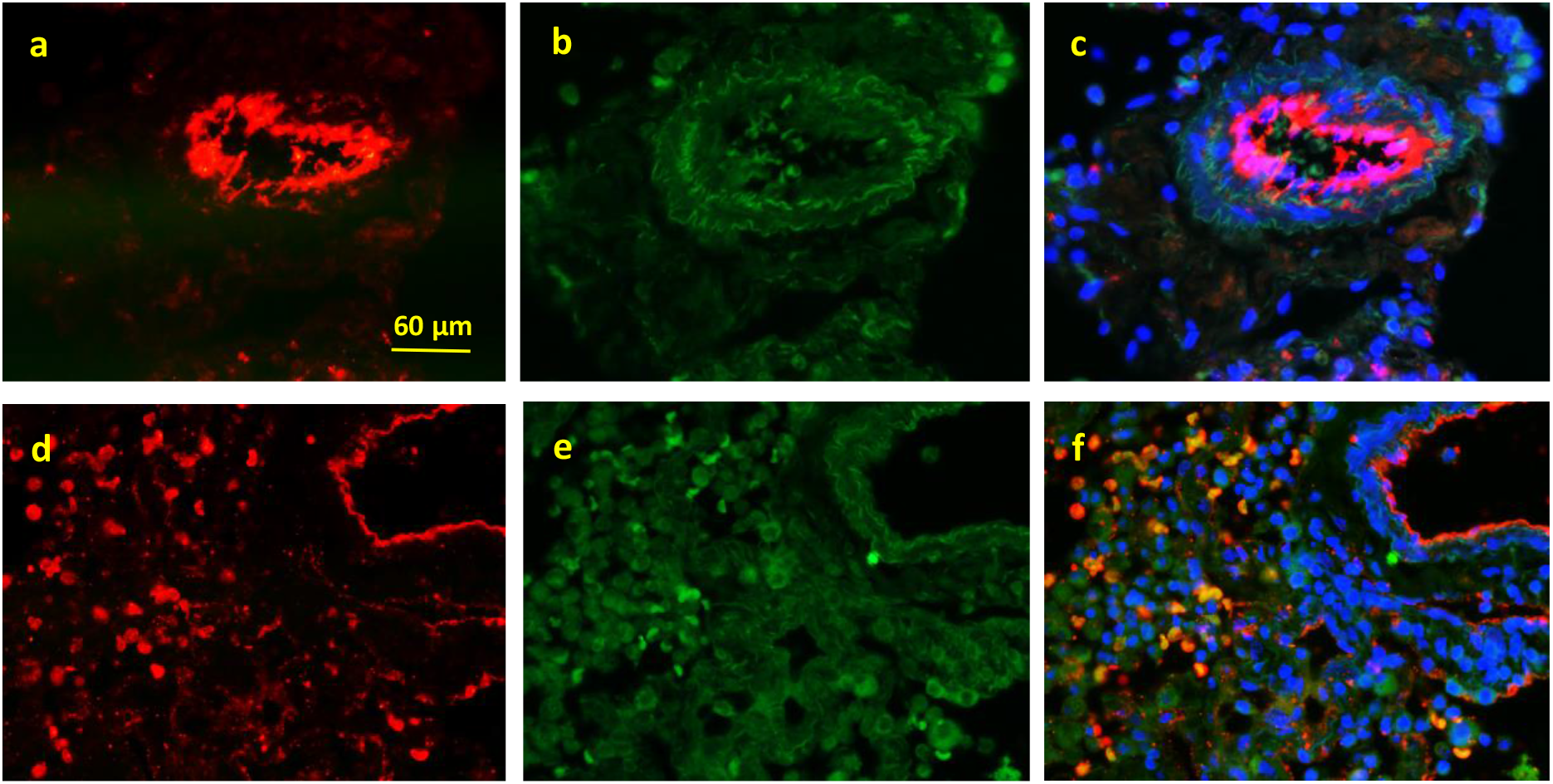
Immuno-staining of vWF and TGF-β on sections of CF and non-CF lungs. (a, b, c), lung sections of non-CF donor; (d, e, f), lung sections of CF donor. (a, d), vWF immune-staining; (b, e), TGF-β immune-staining; (c, f), merged images. All the pictures were taken under the same setting (bar = 60 µm).

To our surprise, the staining pattern of vWF was completely different in CF lungs when compared to that of non-CF lungs: in addition to the conventional localization in the vessel-like structure (Figure 9 a, d), a strong vWF signal was detected in individual and disk-shaped cells (Figure 9d). Moreover, TGF-β and vWF were co-localized in these cells (Figure 9 f).

To confirm these results of vWF mislocalization, we performed immuno-staining on purified RSC cells cultured *in vitro*. Again, we observed a uniform and strong signal of vWF on the cytoplasmic membrane of CF RSC cells (Figure 10 c, e). In contrast, vWF-staining in non-CF RSC cells appeared as dots (Figure 10 a). Therefore, the vWF was indeed mislocalized as well as over-expressed in CF RSC cells. Consistent with our observation, it has been reported that the circulating level of vWF, t-PA and P-selectin was significantly higher in CF patients than in non-CF patients [15]. As far as we know, this is probably the first example of cytoplasmic membrane localization of vWF that is normally stored in Weibel-Palate bodies or secreted. This could explain why the CF RSC cells looked bigger when labeled by anti-vWF antibody than by anti-TGF-β antibody, since it is well-documented that vWF forms ultra-large polymers [5]. We believe that the mislocalization of vWF, together with TGF-β expression in RSC cells, may have caused the lung fibrosis and inflammation in CF patients, even in absence of viral and bacterial infections. Thus, this is a cell autonomous phenotype of CFTR mutations. However, it’s unclear how defective CFTR function led to mislocalization of vWF in CF RSC cells.

**Figure 10.**
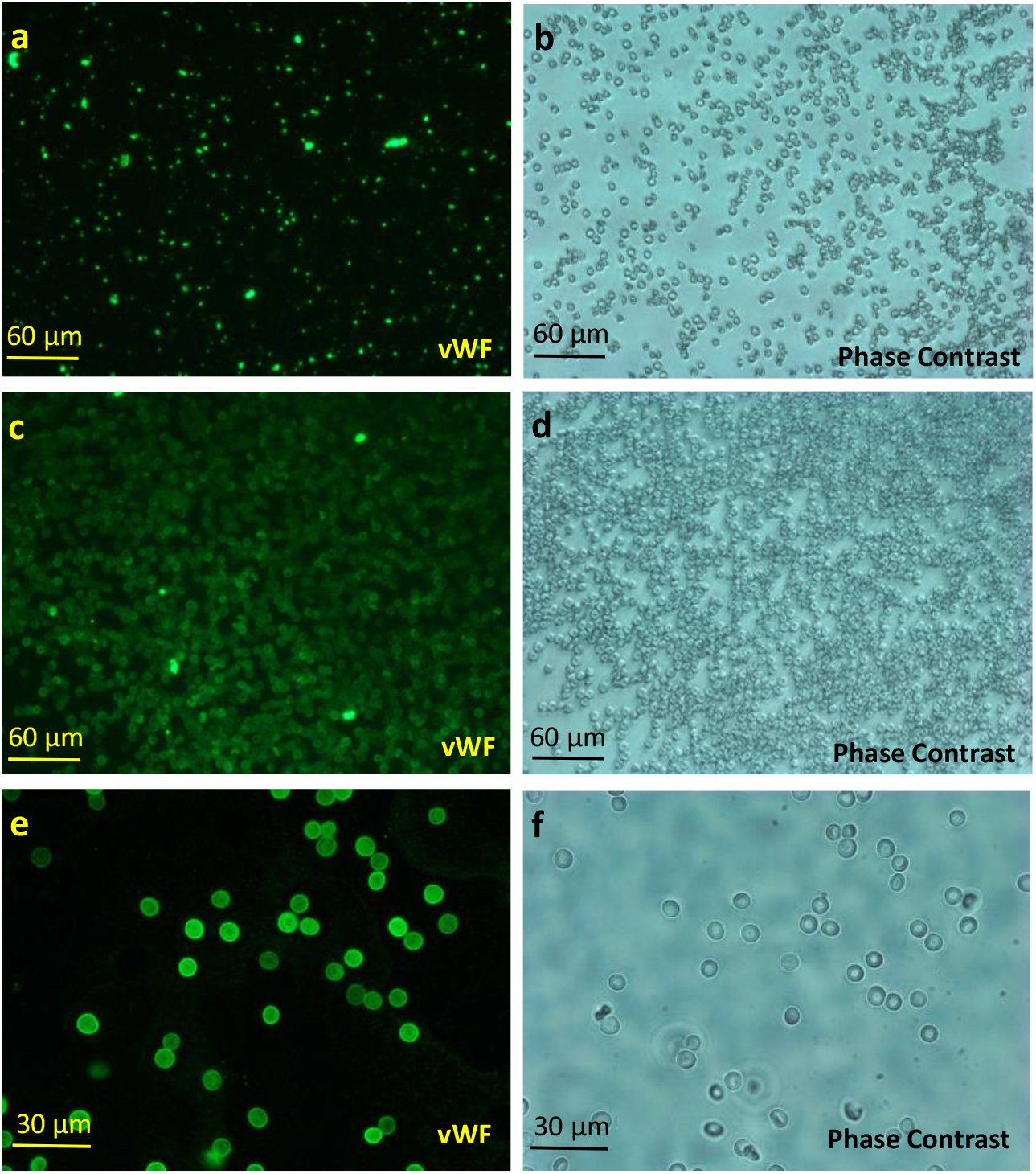
Immuno-staining of RSC cells isolated from CF and non-CF lungs cultured *in vitro*, with anti-vWF antibody. (a, c, e), Immuno-staining for vWF. (b, d, f), phase contrast images. (a, b), RSC cells from non-CF donor. (c, d. e, f), RSC cells from CF donor.

### Discussion and conclusion

When we see something red in human or in animal bodies or biopsies, we would immediately conclude that it must be blood, more precisely the red blood cells (erythrocytes). The association between the red color and the red blood cells is so strong that it leaves no room for other interpretations. In order to reconstitute the airway epithelial models *in vitro*, we isolate routinely the epithelial cells from all kinds of biopsies from respiratory system, such as nasal polyps, pieces of bronchus, or lung parenchyma, or the whole lungs. Some red colored cells are always present, and sometimes in large quantities. However, the presence of these red cells didn’t bother us, in both literal and figurative senses of the term. In fact, these red cells, except forming aggregates or rouleaux, are not very active in culture, and they disappeared from the culture dishes with time. But we were still intrigued by the presence of large quantities of red cells in the biopsies from the Cystic Fibrosis patients. In fact, the lung parenchyma of CF patients was filled with red cells. If they were Red Blood Cells, this means that most of the CF patients would have had severe hemorrhages in the lungs, and they may have died from bleeding. However, bleeding is rather rare in CF patients. The main cause of mortality of CF patients is respiratory failure due to airway obstruction and chronic infections [9, 10, 11, 12]. Thus, what are these red cells? In this article, we try to demonstrate that the red cells present in the biopsies are not the red blood cells, rather a population of red cells located outside of the blood vessels, which may have multiple and important functions.

First, with Dapi staining, we showed that these red cells have a nucleus. This feature alone would be sufficient to rule out the possibility of being red blood cells. Another characteristic of RBCs is their mobility: in order to perform their functions, RBCs have to circulate constantly in the blood vessels. As we mentioned in the “Introduction”, in a healthy person, the blood as well as RBC is confined within the cardiovascular system, which in the lungs has a tree-like hierarchical branching structure. By injecting a blue ink into the right ventricle of the rat lungs, we could label the vessels in blue color, which could be clearly seen under contrast microscope in intact lung and in cleared lung slices. Outside of the blue vessels, a large amount of red colored cells could be observed alongside of both arterioles and capillary bed. The same was true for human lungs. These red cells formed integral part of the lung tissues, either individually or collective as densely packed clusters.

We provided also some evidence suggesting that these red cells are part of the connective tissues of the lung. First, we observed that they attached to and intermingled with the collagen fibers in the biopsies. Second, they could be isolated more easily with collagenase than with other enzymes. Third, they attached spontaneously to and formed aggregates on collagen fibers *in vitro*, displaying a collaborative and collective behavior.

Consistent with these observations, Red soma cells expressed vWF, a large multimeric glycoprotein present in blood plasma and produced constitutively as ultra-large vWF polymers in endothelium, megakaryocytes, and sub-endothelial connective tissue. vWF mediates the adhesion of platelets to sites of vascular damage by binding to specific platelet membrane glycoproteins and to constituents of exposed connective tissue [5]. Since RSC cells expressed vWF and PF4, we wondered if RSC cells may also play a role in coagulation. Interesting, upon exposure to toxic chemicals, PSS for example, RSC cells were rapidly (within minutes) activated and transformed into filaments and formed dense networks. If this was not an artefact that only produces *in vitro* after being exposed to toxic chemicals, then this way of coagulation would be much more efficient than that to recruit platelets from the blood, because of their large number, high cell density as well as their close location to blood vessels.

One of the important factors in coagulation is vWF. Interesting, we observed an abnormal localization and an over-expression of vWF in RSC cells derived from CF patients, both on the lung sections and in individual RSCs cultured *in vitro*. Unlike the staining pattern of RSCs from normal donors, all the RSC cells from CF donors were stained positive by anti-vWF. Furthermore, the same Red Soma Cell looked much bigger when labeled by anti-vWF antibody than that by anti-TGF-β antibody. These results suggest that, instead of being secreted, the vWF in CF RSC cells was probably tethered on the cytoplasmic membranes. As far as we know, this phenomenon has not been observed before. Since vWF binds specifically to several collagens *in vitro*, including types I, II, III, IV, V, and VI [5], the membrane bound-vWF may enhance the attachment of the CF-RSC to collagen fibers, and also the ability of CF-RSC cells to form dense clusters. Thus, the mislocalization of vWF, together with TGF-β expression, could be the molecular basis of the formation of big lumps of red tissues surrounding the bronchi of CF donors, and the fibrotic nature of the CF lung parenchyma.

So what are these RSC cells? If we take an analytical approach that we are used to, these red cells, judging by their color and morphology, could indeed be considered as red blood cells. Based on the biomarkers, they could be classified as megakaryocytes/platelets, because they express PF4; or as endothelial cells, due to the presence of vWF; or as alveolar macrophages, since they are positive for SCF-1R, CD200R, TGF-β, etc… Clearly, such reasoning is wrong. These cells may have the potential to become any of these cells, but they are none of them. Thus, the use of any existing nomenclature to describe these red cells would inevitably lead to confusion, and misunderstanding. In order to comprehend fully the nature and function of this population of red cells, a holistic approach would be needed. We should consider these red cells as a new and single cell type, with well-defined characteristics and functions. No doubt, much more research efforts would be needed to fully understand these cells. But, to begin with, it would be sufficient to call this population of red cells with a new name, Red Soma Cells, to distinguish these cells from the Red Blood Cells,

Not surprising, such cells have been observed and described from different angles previously. Just to cite a few examples here:

1. Swirski et al. reported that in the spleen of mouse, there exists a population of resident bona fide undifferentiated monocytes, outnumbering their equivalents in circulation. The reservoir monocytes assemble in clusters in the cords of the subcapsular red pulp and are distinct from macrophages and DCs. In response to ischemic myocardial injury, splenic monocytes increase their motility, exit the spleen en masse, accumulate in injured tissue, and participate in wound healing [16].
2. Lefrançais et al. demonstrated that the mouse lung is a major site of platelet biogenesis as well as a reservoir for hematopoietic progenitors [17]. By transplanting profused mouse lungs into thrombocytopenic animals, deficient in platelet and hematopoietic stem cells, the authors discovered that lung hematopoietic progenitors are able to migrate to the bone marrow, reconstitute normal platelet counts, and contribute to the hematopoiesis of several cell lines several months after transplantation. In our opinion, the monocytes in the mouse spleen described by Swirski et al., the hematopoietic progenitors in the mouse lung implied by Lefrançais et al., and the Red Soma Cells described here are the same cells. Therefore, based on all these findings, we would like to propose a general concept: What makes our body look red is not the Red Blood Cells, rather the Red Soma Cells. RSC cells are stem cells, located as clusters in the connective tissues of our body rich in collagen fibers. They function as a unit, displaying collective and collaborative behaviors.

Our assumption seems to be a leap of faith. Indeed, we have no experimental data on other organs or parts of our body. However, it shouldn’t be forgotten that our body is an interconnected whole. If monocytes can exit spleen and travel all the way to the injured heart, and hematopoietic progenitors in the lungs can go “back” to the irradiated bone marrow, we don’t see any obstacle for RSC cells to move around and colonize all the connective tissues in our body. Judging by their large number and rapid mobilization upon injury, we believe the RSC cells function as our “bodyguard”, literally and figuratively. This may be the reason why RSC cells exist as densely packed clusters. They may also be involved various inflammatory diseases, such as Cystic fibrosis. In addition to generate other types of cells, we speculate that they may also participate directly as effectors to fight the invading pathogens by aggregating physically on pathogens, or via release of certain anti-viral or anti-bacterial reagents. Thus, these Red Soma Cells can be considered as an **emergency system** which can be activated rapidly and massively in case of injury, or upon viral and bacterial infections. If true, the red blisters on the skin of people with Chickenpox may be a sign of their presence during viral infection. It would not be surprising to find out that RSC cells may also be implicated in pathogenesis of COVID-19, since almost all the stem cells express ACE-2 proteins on their cell surface and respond to the Spike protein of SARS-CoV-2 [18, 19].

In conclusion, what makes our body look red is not the Red Blood Cells, rather the Red Soma Cells. Our discovery of Red Soma Cells will have important theoretic as well as clinical implications.

## Acknowledgements

We thank all the members of Epithelix’ team for their supports. In particular, Samuel Constant and Ludovic Wiszniewski for financial support and scientific discussions. Professor Edouard Sage for providing us the CF biopsies (Service de Chirurgie Thoracique et Transplantation Pulmonaire, Hôpital Foch, 40 rue Worth, 92151 Suresnes, France). Song Huang made the discovery, designed and performed most of the experiments, and wrote the article. Cindia Lopez performed the immuno-stainings presented in Figure 4, 6, 9, 10.

## Reference

1. Wikipedia contributors. (2021, February 11). Red blood cell. In Wikipedia, The Free Encyclopedia. Retrieved 09:38, March 6, 2021. from https://en.wikipedia.org/w/index.php?title=Red_blood_cell&oldid=1006158650.

2. Wikipedia contributors. (2021, February 24). Circulatory system. In Wikipedia, The Free Encyclopedia. Retrieved 09:40, March 6, 2021, from https://en.wikipedia.org/w/index.php?title=Circulatory_system&oldid=1008653141.

3. Alberts, B., Johnson, A., Lewis, J., Raff, M., Roberts, K., & Walter, P. (2002). Molecular biology of the cell. New York: Garland Science.

4. Guibert, E. E., Petrenko, A. Y., Balaban, C. L., Somov, A. Y., Rodriguez, J. V., & Fuller, B. J. (2011). Organ Preservation: Current Concepts and New Strategies for the Next Decade. Transfusion medicine and hemotherapy: offizielles Organ der Deutschen Gesellschaft fur Transfusionsmedizin und Immunhamatologie, 38(2), 125–142. https://doi.org/10.1159/000327033.

5. Sadler JE. (1998). Biochemistry and genetics of von Willebrand factor. Annu Rev Biochem. 67: 395–424. doi: 10.1146/annurev.biochem.67.1.395. PMID: 9759493.

6. Newton, JR; Ah-See, KW (April 2008). 舠A review of nasal polyposis”. Therapeutics and Clinical Risk Management. 4 (2): 507–512. doi:10.2147/tcrm.s2379. PMC 2504067. PMID 18728843.

7. DeMuri, Gregory (2015). Mandell, Douglas, and Bennett’s Principles and Practice of Infectious Diseases. pp. 774–784. ISBN 978-0443068393.

8. Wikipedia contributors. (2021, February 24). Root. In Wikipedia, the Free Encyclopedia. Retrieved 10:09, March 6, 2021, from https://en.wikipedia.org/w/index.php?title=Root&oldid=1008615078.

9. Collins, F. S. Cystic Fibrosis: Molecular Biology and Therapeutic Implications. Science 256, 774–779 (1992).

10. Davies, J. C., Alton E. W. F. W. & Bush, A. Cystic fibrosis. BMJ 335, 1255–1259 (2007).

11. Boucher, R. C. New concepts of the pathogenesis of cystic fibrosis lung disease. Eur. Respir. J. 23, 146–158 (2004).

12. Gibson, R. L., Burns, J. L. & Ramsey, B. W. (2003). Pathophysiology and Management of Pulmonary Infections in Cystic Fibrosis. Am. J. Respir. Crit. Care Med. 168, 918–951

13. Yue, X., Shan, B., & Lasky, J. A. (2010). TGF-β: Titan of Lung Fibrogenesis. Current enzyme inhibition 6(2), 10.2174/10067. https://doi.org/10.2174/10067.

14. Aschner, Y., & Downey, G. P. (2016). Transforming Growth Factor-β: Master Regulator of the Respiratory System in Health and Disease. American journal of respiratory cell and molecular biology 54(5), 647–655. https://doi.org/10.1165/rcmb.2015-0391TR.

15. Romano M, Collura M, Lapichino L, Pardo F, Falco A, Chiesa PL, Caimi G, DavÌ G. (2001). Endothelial perturbation in cystic fibrosis. Thromb Haemost. 86(6):1363–7. PMID: 11776300.

16. Swirski FK, Nahrendorf M, Etzrodt M, Wildgruber M, Cortez-Retamozo V, Panizzi P, Figueiredo JL, Kohler RH, Chudnovskiy A, Waterman P, Aikawa E, Mempel TR, Libby P, Weissleder R, Pittet MJ. (2009). Identification of splenic reservoir monocytes and their deployment to inflammatory sites. Science 325(5940):612–6. doi: 10.1126/science.1175202. PMID: 19644120; PMCID: PMC2803111.

17. Lefrançais E, Ortiz-Muñoz G, Caudrillier A, Mallavia B, Liu F, Sayah DM, Thornton EE, Headley MB, David T, Coughlin SR, Krummel MF, Leavitt AD, Passegué E, Looney MR. (2017) The lung is a site of platelet biogenesis and a reservoir for hematopoietic progenitors. Nature 544(7648):105–109. doi: 10.1038/nature21706. Epub 2017 Mar 22. PMID: 28329764; PMCID: PMC5663284.

18. Zambidis, E. T., Park, T. S., Yu, W., Tam, A., Levine, M., Yuan, X., Pryzhkova, M., & Péault, B. (2008). Expression of angiotensin-converting enzyme (CD143) identifies and regulates primitive hemangioblasts derived from human pluripotent stem cells. Blood 112(9), 3601– 3614. https://doi.org/10.1182/blood-2008-03-144766.

19. Mariusz Z. Ratajczak, Kamila Bujko, Andrzej Ciechanowicz, Kasia Sielatycka, Monika Cymer, Wojciech Marlicz, Magda Kucia (2021) SARS-CoV-2 Entry Receptor ACE2 Is Expressed on Very Small CD45− Precursors of Hematopoietic and Endothelial Cells and in Response to Virus Spike Protein Activates the Nlrp3 Inflammasome. Stem Cell Rev Rep. 17(1): 266–277. Published online 2020 Jul 20. doi: 10.1007/s12015-020-10010-z PMCID: PMC7370872

